# Relaxed constraint and functional divergence of the progesterone receptor (PGR) in the human stem-lineage

**DOI:** 10.1101/799569

**Authors:** Mirna Marinić, Vincent J. Lynch

## Abstract

The steroid hormone progesterone, acting through the progesterone receptor (PR), a ligand-activated DNA-binding transcription factor, plays an essential role in regulating nearly every aspect of female reproductive biology. While many reproductive traits regulated by PR are conserved in mammals, Catarrhine primates evolved several derived traits including spontaneous decidualization, menstruation, and a divergent (and unknown) parturition signal, suggesting that PR may also have evolved divergent functions in Catarrhines. There is conflicting evidence, however, whether the progesterone receptor gene (*PGR*) was positively selected in the human lineage. Here we show that *PGR* evolved rapidly in the human stem-lineage (as well as other Catarrhine primates), which likely reflects an episode of relaxed selection intensity rather than positive selection. Coincident with the episode of relaxed selection intensity, ancestral sequence resurrection and functional tests indicate that the major human PR isoforms (PR-A and PR-B) evolved divergent functions in the human stem-lineage. These results suggest that the regulation of progesterone signaling by PR-A and PR-B may also have diverged in the human lineage and that non-human animal models of progesterone signaling may not faithfully recapitulate human biology.

## Introduction

The steroid hormone progesterone plays a central role in female reproduction. In Eutherian (Placental) mammals, for example, progesterone regulates the timing of the reproductive cycle, ovulation, decidualization, implantation and the maintenance of pregnancy, myometrial quiescence, and the cessation of pregnancy at parturition (Lydon et al. 1995; Conneely et al. 2001; Kubota et al. 2016). Many of the biological actions of progesterone are mediated through the progesterone receptor (PR), which acts as a ligand-activated DNA-binding transcription factor (Lydon et al. 1995; Conneely et al. 2001; Wetendorf and DeMayo 2012). The progesterone receptor gene (*PGR*) encodes two well-characterized isoforms (PR-A and PR-B) that are transcribed from distinct promoters and utilize different translation start sites in the first exon, but are identical except for a 165 amino acid trans-activation domain in the amino terminus of PR-B (Kastner et al. 1990). Consistent with the structural differences between PR-A and PR-B, previous studies have shown that PR-B is a stronger trans-activator of progesterone responsive genes than PR-A, whereas PR-A acts as a dominant trans-repressor of PR-B mediated trans-activation (Kastner et al. 1990; Huse et al. 1998; Giangrande et al. 2000; Abdel-Hafiz et al. 2002; Kaya et al. 2015). These opposing transcriptional activities result from different post-translational modifications and cofactor interactions. For example, PR-A does not efficiently interact with co-activators but strongly binds the co-repressor SMRT, allowing it to function as a dominant trans-repressor (Huse et al. 1998; Giangrande et al. 2000; Abdel-Hafiz et al. 2002).

While female reproductive traits, particularly those regulated by PR, are generally well conserved across Eutherian (Lombardi 1998) and Therian mammals (Renfree and Shaw 2001; Behringer et al. 2006; Bradshaw and Bradshaw 2011), the genes that underlie them can evolve rapidly, often under the influence of positive selection (Arbeitman et al. 2002; Meiklejohn et al. 2003; Parisi et al. 2003). Two primate lineages differ dramatically in some otherwise conserved female reproductive traits. Catarrhine primates have evolved spontaneous differentiation (decidualization) of endometrial stromal fibroblasts (ESFs) into decidual stromal cells (DSCs) under the direct action of progesterone (Mess and Carter 2006; Kin et al. 2014), menstruation (Emera et al. 2012; Strassmann 2015), and a divergent (and unknown) parturition signal (Csapo 1956; Csapo 1965). In addition, humans have evolved permanently enlarged breasts (Hamilton 1984; Mascia-Lees et al. 1986), concealed ovulation (Benshoof and Thornhill 1979; Burley 1979), and longer pregnancy and labor compared to other primates (Graham 1981; Chen et al. 2008; Phillips et al. 2015). These data suggest that PR may have evolved rapidly in humans and Catarrhines.

There is conflicting evidence whether *PGR* was positively selected in the human lineage. *PGR* was ranked 8^th^ in a genome-wide scan for positive selection in human-chimp-mouse gene trios (Clark et al. 2003) and 29^th^ among genes with the strongest statistical evidence of positive selection in a pair-wise genome-wide scan for positively selected genes in human and chimpanzee genomes (Nielsen et al. 2005). In contrast, evidence of positive selection on *PGR* was not detected in the human lineage in analyses of human-chimp-macaque gene trios (Bakewell et al. 2007), human-chimp-mouse-rat-dog orthologs (Arbiza et al. 2006), a dataset including seven primates (George et al. 2011), or a dataset including nine primates (van der Lee et al. 2017). A study of human-chimp-macaque-mouse-rat-dog orthologs found evidence for a positively selected class of sites in the human lineage (4.45%, *d*_*N*_/*d*_*S*_ =3.15), but the results were not statistically significant (LRT *P*=0.46) (Kosiol et al. 2008). The most comprehensive analysis of *PGR* evolution thus far, which included 14 primates and four outgroups (Chen et al. 2008), found evidence for an episode of rapid evolution in the human lineage. However, this study did not explicitly test whether the *d*_*N*_/*d*_*S*_ rate was significantly greater than 1, and while a branch-sites test identified a proportion of sites as potentially positively selected (17%, *d*_*N*_/*d*_*S*_ =5.5), the null hypothesis could not be rejected (LRT *P*=0.22).

To resolve these conflicting data, we assembled a dataset of 119 Eutherian *PGRs*, including species and sub-species from each primate lineage, as well as modern and archaic humans (Neanderthal and Denisovan), and used a suite of maximum likelihood-based methods to characterize the strength and direction of selection acting on *PGR*. We found that *PGR* evolved rapidly in the human lineage, however, there was little evidence it was driven by an episode of positive selection. Rather, the rate acceleration was consistent with a relaxation in the intensity of purifying selection, which likely reflects a long-term trend in Catarrhine primates. To test whether the episode of relaxed constraint occurred coincident with a change in the function of the human PR, we resurrected the ancestral human (AncHuman) and human/chimp (AncHominini) PR-A and PR-B isoforms and tested their ability to trans-activate reporter gene expression from the decidual *prolactin* promoter (dPRL-332), a well-characterized progesterone responsive element (Pohnke et al. 1999; Christian, Pohnke, et al. 2002; Christian, Zhang, et al. 2002; Lynch et al. 2009; Jiang et al. 2011). We found pronounced functional differences between AncHuman and AncHominini PR isoforms, particularly in the ability of PR-A to trans-repress PR-B. These data suggest that an episode of relaxed purifying selection altered the function of the human PR, which may have impacted female reproductive biology.

## Results

### No evidence of positive selection in the human stem-lineage

We assembled a dataset of 119 Eutherian *PGRs*, including representatives of each major primate lineage, to characterize the strength and direction of selection generally acting on these genes and explicitly test for an episode of positive selection in the human lineage (**Figure 1**). We first compared two likelihood models using HyPhy (Pond et al. 2005): one in which *d*_*N*_/*d*_*S*_ ratio (ω) in the human stem-lineage was estimated separately from all other lineages, and a second that had a single ω for all lineages. We allowed for variable *d*_*N*_ and *d*_*S*_ cross sites with 3 rate classes in both models. The two ω model was significantly better than the single ω model, with ω_2_=2.67 (95% CI=1.15-5.18; LRT=8.91, *P*=2.83×10^−3^), however, the *d*_*N*_/*d*_*S*_ ratio in the human stem-lineage was not significantly different than 1 (LRT=1.15, *P*=0.28). To test if there was a class of sites with ω>1 in the human stem-lineage, we used an adaptive branch-site random effects likelihood (aBSREL) model (Kosakovsky Pond et al. 2011) that infers the optimal number of site classes and the ω of each site class for each lineage. The aBSREL model inferred a class of sites in the human stem-lineage with ω=2.80, but ω was not significantly different than 1 (*P*=0.44). We also used the branch-site unrestricted statistical test for episodic diversification (BUSTED) model (Murrell et al. 2015), which can detect positive selection on a subset of branches and at a subset of sites, and again did not infer either episodic diversifying selection (*P*=0.57), or support for positive selection on individual substitutions in the human stem-lineage (evidence ratios ~2). In contrast, aBSREL and BUSTED did identify evidence of positive selection in non-primate lineages (**Supplementary Tables**).

**Figure 1.**
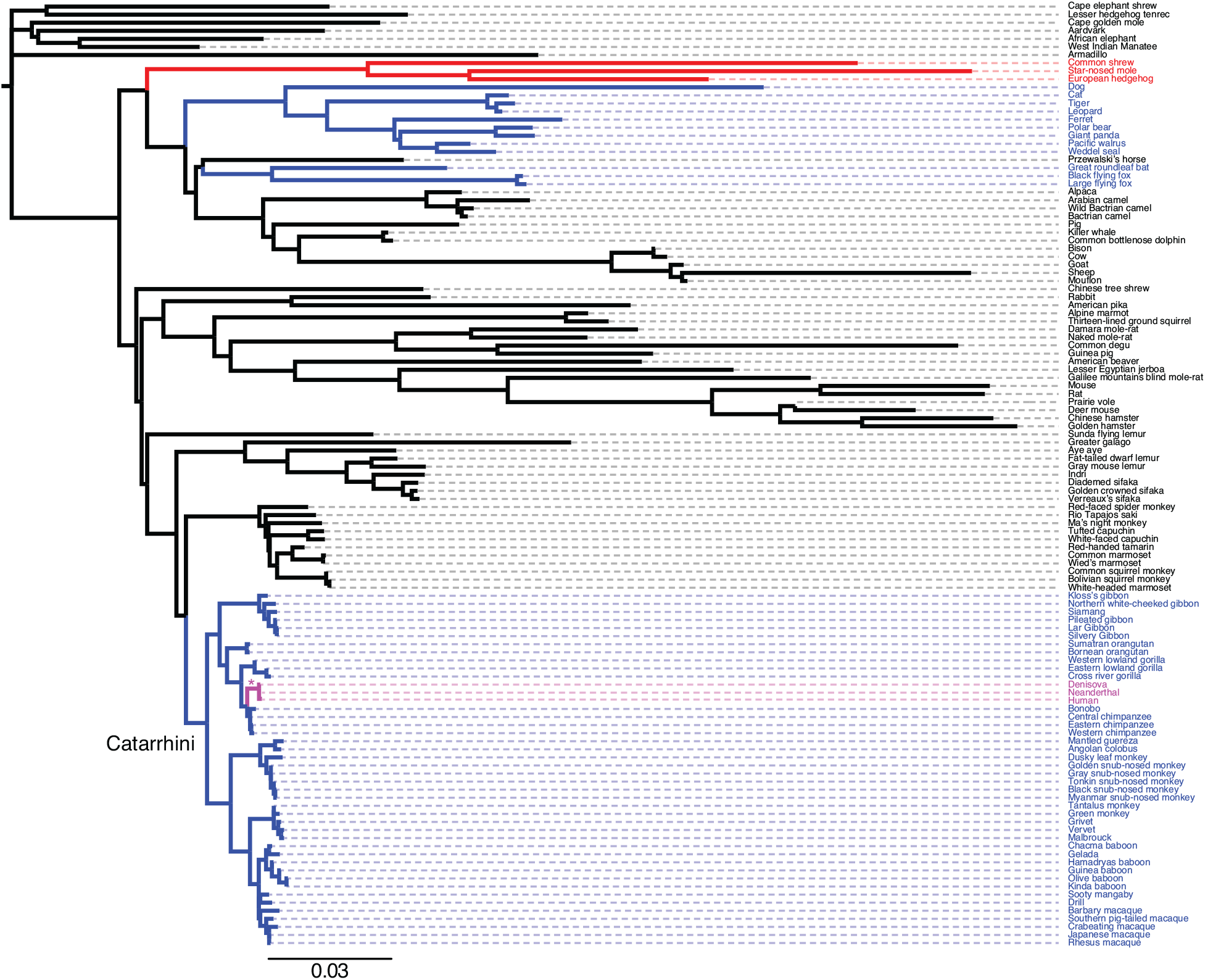
Phylogeny of species used to characterize the strength and direction of selection on *PGR* genes. Branch lengths are drawn proportional to the number of substitutions per site under the GTR model. The human stem-lineage, modern human, Denisovan, and Neanderthal lineages are shown in magenta. Asterisk (*) indicates the human stem-lineage, which has an ω=1.06 and k=0.00 (LRT=5.01, *P*=0.01) under the RELAX *a priori* model. Branches are colored black if k is not significantly different than 1, red if k is significantly greater than 1 and blue if k is significantly less than 1 in the *ad hoc* RELAX analyses.

### No evidence of positive selection at sites with human-specific substitutions

The methods used above can detect positive selection acting on *a priori* defined lineages (two ω-ratio model) and classes of sites across all lineages (aBSREL and BUSTED), but cannot detect positive selection acting on specific sites, alignment-wide. To test for pervasive positive and negative selection at individual amino acid sites, we used two distinct models: Fixed Effects Likelihood (FEL) and Single-Likelihood Ancestor Counting (SLAC) (Pond and Frost 2005). FEL, which uses a maximum-likelihood approach to infer *d*_*N*_ and *d*_*S*_ substitution rates on a per-site basis, found seven sites with evidence of pervasive positive selection and 583 sites with evidence of purifying selection (**Supplementary Tables**). Similarly, the SLAC model, which combines maximum-likelihood and counting approaches to infer *d*_*N*_ and *d*_*S*_, showed evidence for positive selection at four sites and purifying selection at 531 sites (**Supplementary Tables**).

Both FEL and SLAC assume that the selection pressure for each site is constant across the entire phylogeny, thus sites are not allowed to switch between rate categories (positive, negative and neutral). This assumption, however, may not be realistic when evolution is episodic rather than pervasive. Therefore, we use the Fast Unconstrained Bayesian AppRoximation (FUBAR) method (Murrell et al. 2013) and the Mixed Effects Model of Evolution (MEME) method (Murrell et al. 2012) to test for episodic positive selection at amino acid sites. In addition to detecting episodic selection, FUBAR may have more power than FEL when positive selection is present but weak (i.e., low values of ω>1). FUBAR identified a single site with evidence of positive selection and 689 sites with evidence of purifying selection (**Supplementary Tables**), whereas MEME found evidence of positive selection at 17 sites (**Supplementary Tables**). Of the eight human-specific substitutions (see below), all occurred at sites that are under negative selection or that evolve neutrally, alignment-wide (**Tables 1–4**). These data suggest that while *PGR* may have experienced positive selection at some sites in some Eutherian lineages, the episode of rapid evolution in the human stem-lineage is unlikely to result from either pervasive or episodic positive selection.

**Table 1.** Fixed Effects Model (FEL) results for sites with human-specific amino acid substitutions. α, synonymous substitution rate. β, non-synonymous substitution rate. ω, *d*_*N*_/*d*_*S*_ ratio. α = β, rate estimate under the neutral model. LRT, likelihood ratio test statistic for α = β versus α < β. Branch length, the total length of branches contributing to the inference at this site (used to scale *d*_*N*_ and *d*_*S*_).

| Site | $\alpha$ | $\beta$ | $\omega$ | $\alpha = \beta$ | LRT | P-value | Branch length |
| --- | --- | --- | --- | --- | --- | --- | --- |
| 159 | 1.42 | 0.63 | 0.44 | 0.85 | 2.80 | 0.10 | 7.37 |
| 172 | 0.86 | 0.66 | 0.77 | 0.72 | 0.24 | 0.62 | 6.18 |
| 192 | 0.27 | 0.38 | 1.40 | 0.35 | 0.17 | 0.68 | 3.02 |
| 201 | 0.54 | 0.30 | 0.56 | 0.38 | 0.69 | 0.41 | 3.21 |
| 216 | 5.15 | 0.21 | 0.04 | 0.63 | 12.36 | 0.00 | 13.97 |
| 347 | 2.76 | 0.25 | 0.09 | 0.86 | 16.41 | 0.00 | 8.33 |
| 430 | 1.24 | 0.41 | 0.33 | 0.64 | 3.11 | 0.08 | 5.58 |
| 662 | 0.14 | 0.43 | 3.08 | 0.35 | 1.40 | 0.24 | 2.98 |

**Table 2.** Single-Likelihood Ancestor Counting (SLAC) results for sites with human-specific amino acid substitutions. ES, expected synonymous sites. EN, expected non-synonymous sites. S, inferred synonymous substitutions. N, inferred non-synonymous substitutions. P[S], expected proportion of synonymous sites. *d*_*S*_, inferred synonymous substitution rate. *d_N_,* inferred non-synonymous substitution rate. P [*d*_*N*_/*d*_*S*_ > 1], binomial probability that S is no greater than the observed value, with P_s_ probability of success. P [*d*_*N*_/*d*_*S*_ < 1], binomial probability that S is no less than the observed value, with P_s_ probability of success. Total branch length, the total length of branches contributing to the inference at this site (used to scale *d*_*N*_ and *d*_*S*_).

| Site | ES | EN | S | N | P[S] | $d_S$ | $d_N$ | $d_N/d_S$ | P [$d_N/d_S > 1$] | P [$d_N/d_S < 1$] | Total branch length |
| --- | --- | --- | --- | --- | --- | --- | --- | --- | --- | --- | --- |
| 159 | 0.97 | 1.99 | 9 | 10 | 0.33 | 9.29 | 5.01 | 0.54 | 0.94 | 0.13 | 3.71 |
| 172 | 0.95 | 1.92 | 6 | 10 | 0.33 | 6.31 | 5.21 | 0.83 | 0.74 | 0.45 | 3.85 |
| 192 | 0.95 | 1.97 | 2 | 6 | 0.33 | 2.10 | 3.04 | 1.45 | 0.49 | 0.79 | 3.76 |
| 201 | 0.98 | 1.94 | 4 | 5 | 0.33 | 4.10 | 2.57 | 0.63 | 0.85 | 0.35 | 3.76 |
| 216 | 1.00 | 2.00 | 4 | 2 | 0.33 | 3.99 | 1.00 | 0.25 | 0.98 | 0.10 | 3.67 |
| 347 | 0.87 | 1.68 | 11 | 3 | 0.34 | 12.60 | 1.79 | 0.14 | 1.00 | 0.00 | 3.59 |
| 430 | 0.90 | 1.84 | 6 | 5 | 0.33 | 6.64 | 2.71 | 0.41 | 0.96 | 0.12 | 3.46 |
| 662 | 0.79 | 2.22 | 1 | 8 | 0.26 | 1.27 | 3.60 | 2.83 | 0.27 | 0.93 | 3.88 |

**Table 3.** Mixed Effects Model of Evolution (MEME) results for sites with human-specific amino acid substitutions. α, synonymous substitution rate. β-, non-synonymous substitution rate for the negative/neutral evolution component. p-, mixture distribution weight allocated to β- (i.e., the proportion of the tree neutrally of or under negative selection). β+, non-synonymous substitution rate for the positive selection/neutral component. p+, mixture distribution weight allocated to β+ (i.e., the proportion of the tree neutrally of or under positive selection). LRT, likelihood ratio test statistic for episodic diversification (i.e., p+ > 0 and β+ > α). *P*-value, asymptotic *P*-value for episodic diversification (i.e., p+ > 0 and β+ > α). # branches under selection, an estimate for how many branches may been under selection at this site. Total branch length, the total length of branches contributing to the inference at this site (used to scale *d*_*N*_ and *d*_*S*_).

| Site | $\alpha$ | $\beta$ - | $p$- | $\beta$ + | $p$+ | LRT | $P$-value | # branches under selection | Total branch length |
| --- | --- | --- | --- | --- | --- | --- | --- | --- | --- |
| 159 | 1.48 | 1.48 | 0.47 | 0.00 | 0.53 | 0.00 | 0.67 | 0 | 7.97 |
| 172 | 0.86 | 0.79 | 0.67 | 0.4 | 0.33 | 0.00 | 0.67 | 0 | 6.21 |
| 192 | 0.27 | 0.27 | 0.04 | 0.38 | 0.96 | 0.17 | 0.54 | 0 | 3.03 |
| 201 | 0.69 | 0.69 | 0.37 | 0.00 | 0.63 | 0.00 | 0.67 | 0 | 3.3 |
| 216 | 5.82 | 5.82 | 0.06 | 0.00 | 0.94 | 0.00 | 0.67 | 0 | 16.32 |
| 347 | 2.75 | 1.61 | 0.16 | 0.03 | 0.84 | 0.00 | 0.67 | 0 | 8.56 |
| 430 | 1.24 | 0.41 | 0.89 | 0.37 | 0.11 | 0.00 | 0.67 | 0 | 5.58 |
| 662 | 0.14 | 0.14 | 0.00 | 0.43 | 1.00 | 1.40 | 0.25 | 0 | 2.98 |

**Table 4.**
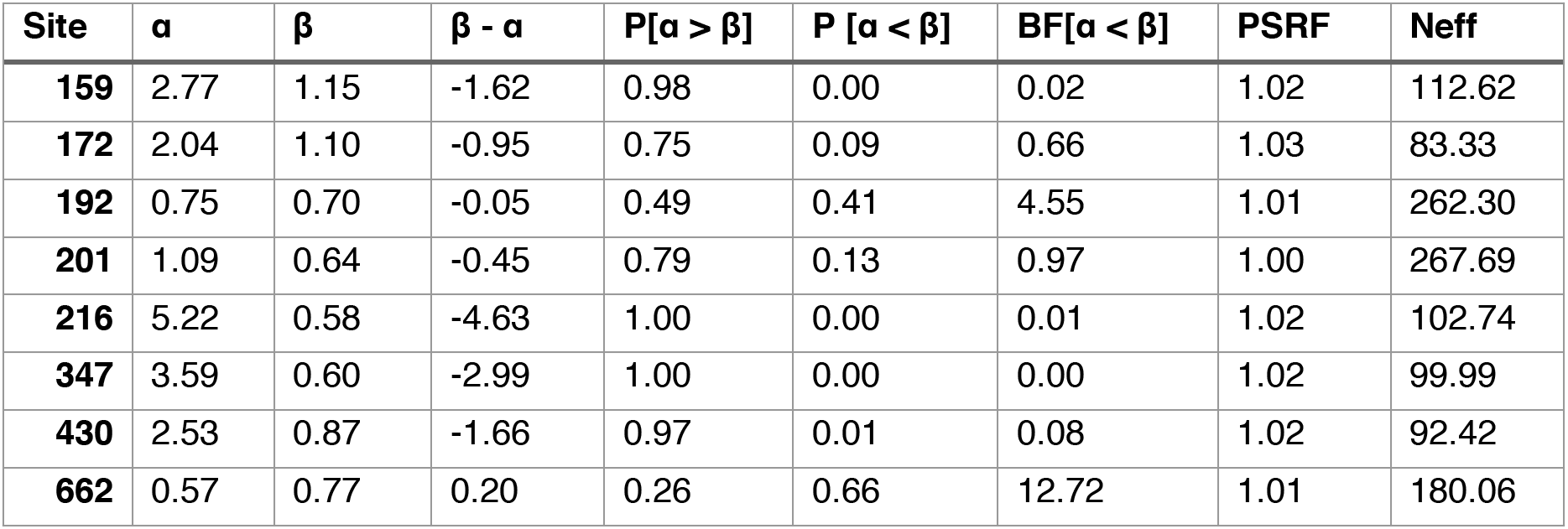
Fast Unconstrained Bayesian AppRoximation (FUBAR) results for site with human-specific amino acid changes. α, mean posterior synonymous substitution rate. β, mean posterior non-synonymous substitution rate. Mean posterior β-α. P[α > β], posterior probability of negative selection. P[α < β], posterior probability of positive selection. BF[α < β], empirical Bayes factor for positive selection. PSRF, potential scale reduction factor (a measure of MCMC mixing). Neff, effective sample size.

### Relaxed purifying selection on human and primate PGRs

To test if the episode of rapid evolution in the human stem-lineage may result from a relaxation in the intensity of selection (both positive and negative), rather than an episode of positive selection, we used a variant of the aBSREL method (RELAX) that explicitly includes a selection intensity parameter (k) (Wertheim et al. 2015). A significant k>1 indicates that selection strength has been intensified, whereas a significant k<1 indicates that the strength of selection (both positive and negative) has been relaxed. RELAX inferred that the baseline rate distribution for branch-site combinations was ω_1_=0.00 (59.41% of sites), ω_2_=0.07 (34.78% of sites), and ω_3_=1.26 (5.81% of sites), while the branch-level relaxation or intensification (k) parameter distribution had a mean of 1.17, median of 0.36, and 95% of the weight within the range of 0.03-6.62. Thus, the ω_3_ class of sites was inferred to be evolving near neutrality (ω=1), alignment-wide. Consistent with an episode of relaxed selection, RELAX inferred ω=1.06 and k=0.00 (LRT=5.01, *P*=0.01) in the human stem-lineage (**Figure 1**).

To explore if the relaxation of constraint was specific to the human lineage or reflects a more widespread pattern, we also tested for relaxed purifying selection in other clades. We found evidence for relaxation in *Homo* (modern and archaic humans; k=0.00, *P*=0.012, LRT=6.38), Hominoidea (apes; k=0.68, *P*=0.09, LRT=2.89), Cercopithecidae (‘Old World monkeys’; k=0.43, *P*=7.28×10^−6^, LRT=20.12), Cercopithecinae (k=0.44, *P*=6.60×10^−4^, LRT=11.60), Colobinae (k=0.32, *P*=1.25×10^−3^, LRT=10.41), and Catarrhini (‘Old World monkeys’ and apes; k=0.43, *P*=1.03×10^−5^, LRT=19.39), but not Platyrrhini (‘New World monkeys’; k=0.95, *P*=0.681, LRT=0.17), Strepsirrhini (lemurs and lorises; k=0.94, *P*=0.37, LRT=0.80), or non-primates (k=2.41, *P*=3.06×10^−8^, LRT=30.67). Thus, we conclude the rate acceleration in the human stem-lineage most likely results from a long-term trend of relaxed purifying selection on Catarrhine primate *PGRs*, rather than an episode of positive selection.

### Human-specific amino acid substitutions are predicted to be deleterious

Similar to previous studies (Chen et al. 2008), we identified eight human-specific amino acid substitutions (G159R, A172V, S192A, G201E, P216A, S347C, P430T, and I662V), six of which are located in the inhibitory function (IF) domain, one in the activation function 3 (AF3) domain, and one in the hinge region (H) (**Figure 2**). While three human-specific substitutions are fixed for the derived amino acid, ancestral variants are segregating at very low frequencies at sites 172 (rs376101426, ancestral allele frequency = 1.37×10^−5^), 201 (rs748082098, ancestral allele frequency = 4.33×10^−6^), 347 (rs11571147, ancestral allele frequency = 9.08×10^−3^), 430 (rs1396844023, ancestral allele frequency = 6.08×10^−6^), and 662 (rs150584881, ancestral allele frequency = 9.59×10^−5^). Similarly, nearly all human-specific amino acid substitutions are fixed for the derived allele in archaic humans, but Neanderthal has 347C whereas Denisovan has 347S (**Figure 2**). We next used Combined Annotation-Dependent Depletion (CADD) (Kircher et al. 2014; Rentzsch et al. 2019) to predict the deleteriousness of each human-specific amino acid substitution and found that most are predicted to be deleterious (**Figure 2**).

**Figure 2.**
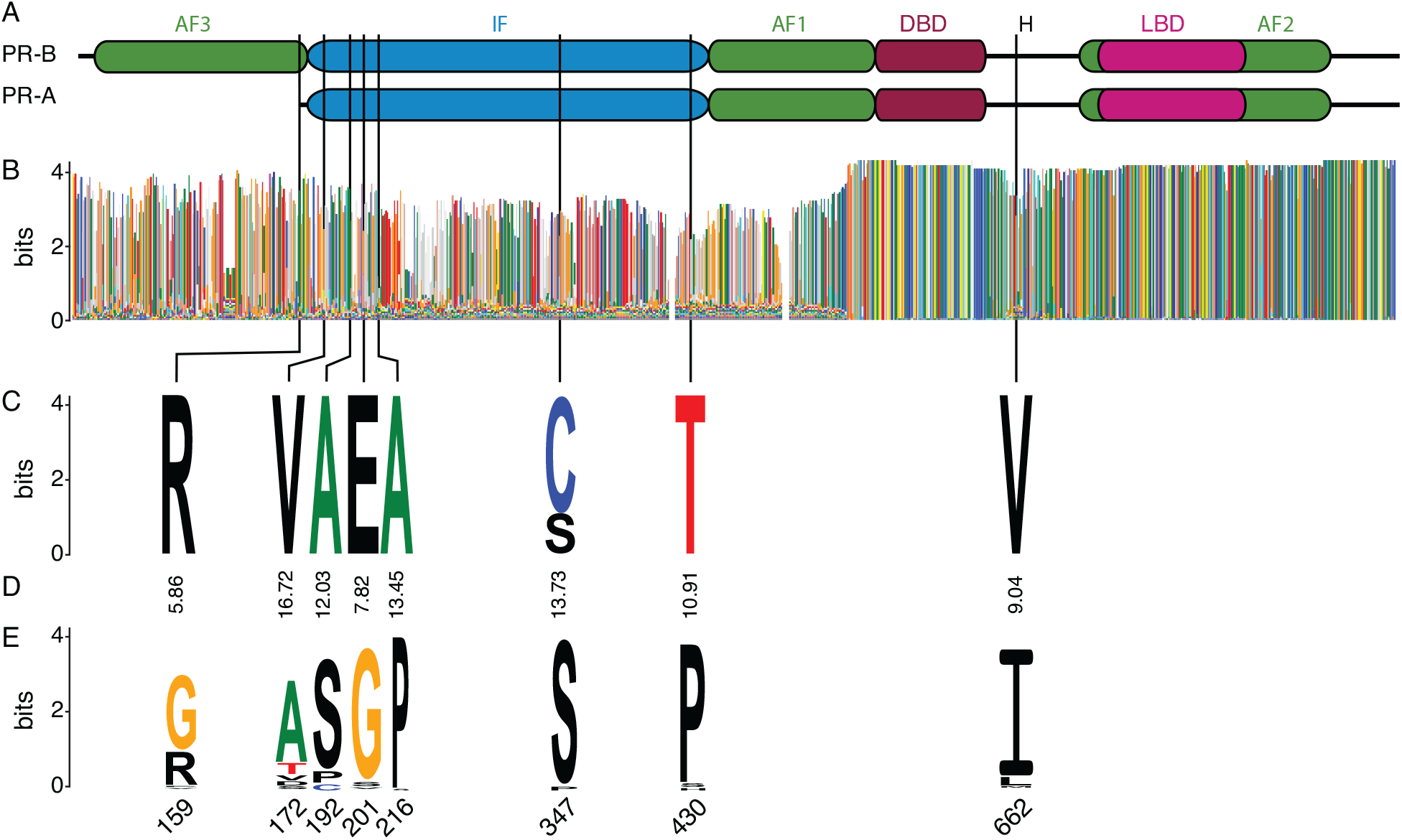
Distribution of human-specific amino acid substitutions in PGR. (A) Functional domains of the progesterone receptor isoforms A (PR-A) and B (PR-B). AF1-3, activation function domains 1-3. IF, inhibitory function domain. DBD, DNA binding domain. LBD, ligand binding domain. (B) Conservation of PGR across 119 Eutherian mammals shown as a logo. Amino acids are colored according to their physical-chemical properties: Neutral amino acids are green, polar amino acids are purple, basic and cysteine are blue, glycine is orange, polar are red, and others are black. (C) Logo showing conservation of human-specific substitutions within modern and archaic humans. Amino acids are colored according to their physical-chemical properties. (D) CADD scores for human-specific substitutions. (E) Logo showing conservation at sites with human-specific substitutions in non-human mammals. Amino acids are colored according to their physical-chemical properties.

### Functional divergence of the human progesterone receptor

To determine if human-specific substitutions have functional consequences, we reconstructed the sequences of the ancestral human (AncHuman) and ancestral human-chimpanzee (AncHominini) PRs (**Figure 3A**), synthesized the PR-A and PR-B isoforms, and cloned them into a mammalian expression vector. Next, we transiently transfected mouse embryonic fibroblasts (MEFs), which do not express *Pgr*, with either AncHuman PR-A, AncHuman PR-B, AncHuman PR-A/PR-B, AncHominini PR-A, AncHominini PR-B, or AncHominini PR-A/PR-B expression vectors and a reporter vector that drives luciferase expression from the decidual *Prolactin* promoter (dPRL-332). The dPRL-332 promoter is a well-characterized progesterone responsive regulatory element that is bound by PR-A and PR-B and directs the expression of *Prolactin* in decidual stromal cells (**Figure 3B**) (Pohnke et al. 1999; Christian, Pohnke, et al. 2002; Christian, Zhang, et al. 2002; Lynch et al. 2009; Jiang et al. 2011). The AncHuman PR-A and AncHominini PR-A isoforms weakly trans-activated luciferase expression from the dPRL-332 promoter, both PR-B isoforms strongly trans-activated luciferase expression, and both PR-A isoforms trans-repressed trans-activated luciferase expression by PR-B (**Figure 3C**). The AncHuman PR-A was a weaker trans-activator than the AncHominini PR-A with a log mean difference of 0.548 (95% CI: 0.099 – 1.03, Mann-Whitney *P*=0.035), while the AncHuman PR-B was a stronger trans-activator than the AncHominini PR-B with a log mean difference of −0.517 (95% CI: −0.779 – −0.276, Mann-Whitney *P*=9.46×10^−4^). Additionally, the ability of AncHuman PR-A to repress AncHuman PR-B was weaker than AncHominini PR-A on AncHominini PR-B with a log mean difference of −1.10 (95% CI: −1.38 – −0.785, Mann-Whitney *P*=3.06×10^−6^) (**Figure 3D**).

**Figure 3.**
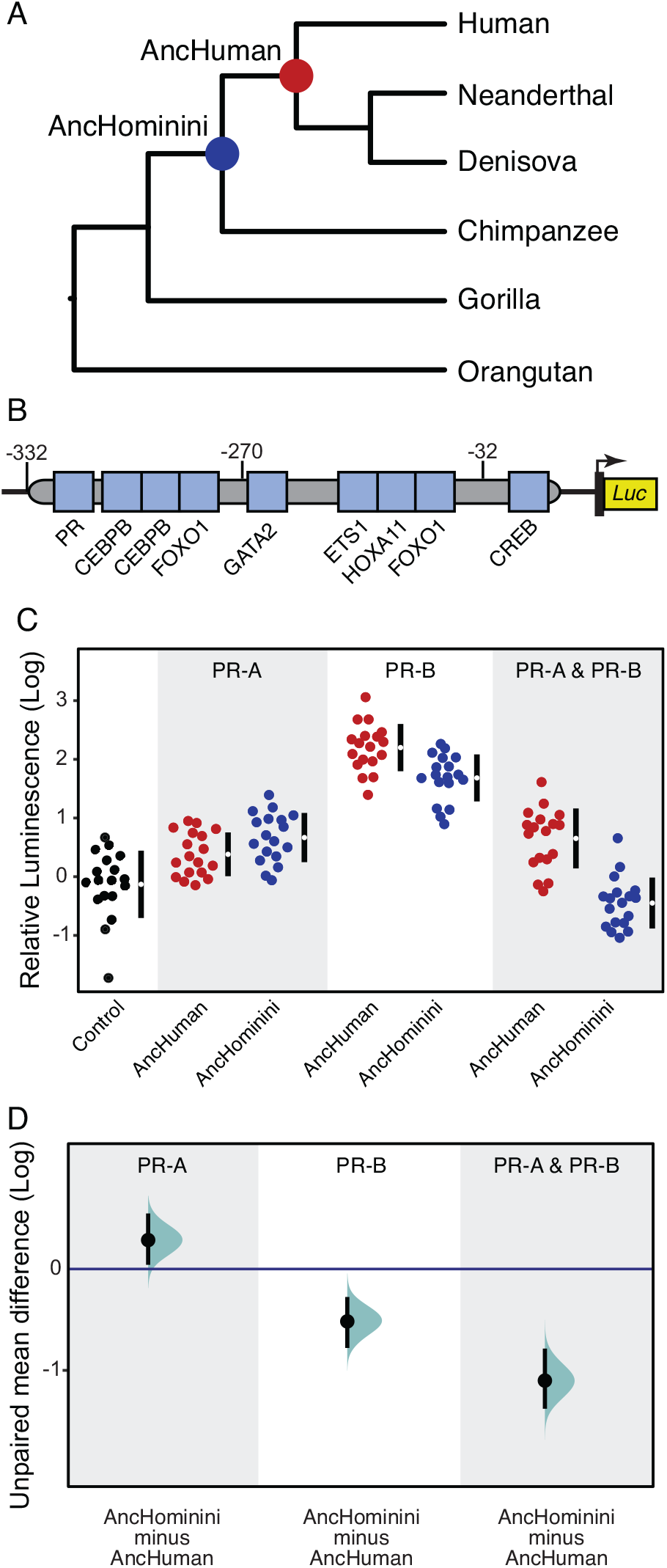
Functional divergence of the human progesterone receptor. (A) Hominin (African ape) phylogeny, nodes with ancestral sequence reconstructions are indicated. (B) The dPRL-332 luciferase reporter vector with the location of experimentally verified transcription factor binding sites. (See main text for references.) (C) Strip chart showing relative luminescence values, standardized to negative control (log scale). Mean is depicted as a white dot; 95% confidence intervals are indicated by the ends of the black vertical error bars. (D) The mean difference in luminescence of AncHuman against AncHominini shown as a Cumming estimation plot (log scale). Mean differences are plotted as bootstrap sampling distributions (*n*=5,000). Each mean difference is depicted as a dot, 95% confidence intervals are indicated by the ends of the black vertical error bars.

## Discussion

### Positive selection or relaxation?

Numerous previous studies have conflicted over whether *PGR* was positively selected in the human lineage (Clark et al. 2003; Nielsen et al. 2005; Arbiza et al. 2006; Bakewell et al. 2007; Chen et al. 2008; George et al. 2011). These studies had limited taxon sampling (particularly within primates) and used models designed to detect positive selection rather than explicitly test evidence for relaxed purifying selection (Wertheim et al. 2015). This may have reduced their power to differentiate an episode of positive selection from a relaxation in the intensity. Our results from multiple methods indicate that *PGR* evolved under relatively strong purifying selection in most lineages and sites and suggest that *PGR* experienced an episode of rapid evolution in the human stem-lineage. However, no method found that an inference of positive selection (ω>1) was supported, whereas several methods found strong support for an episode of relaxed constraint (ω=1) on at least some sites in the human stem-lineage. Our results also suggest that the intensity of purifying selection acting on Catarrhine primate *PGRs* has been relaxed, implying that the episode of relaxed selection intensity in the human stem-lineage reflects a wider trend in Catarrhines.

### Functional divergence of human progesterone receptor

Regardless of the statistical evidence supporting positive selection or relaxed constraint, we found that human PR-A and PR-B have diverged in function compared to the human-chimp ancestor. The trans-activation strength of the AncHuman PR-A is slightly weaker than the AncHominini PR-A, whereas the AncHuman PR-B more strongly trans-activates luciferase expression from the dPRL-332 promoter than the AncHominini PR-B. In addition to acting as a ligand-activated transcriptional activator, PR-A also has the ability to trans-repress the transcriptional activity of PR-B and other nuclear receptors, such as estrogen and glucocorticoid receptors, through its inhibitory function domain (Huse et al. 1998). The most striking difference between the AncHuman and AncHominini isoforms is in the trans-repressive strength of PR-A on PR-B – the AncHominini PR-A can almost completely repress trans-activation by AncHominini PR-B, while the AncHuman PR-A’s capacity to inhibit trans-activation by AncHuman PR-B is weakened compared to AncHominini PR-A. Thus, PR-B can still function as a trans-activator in the presence of stoichiometric ratios (1:1) of PR-A. These results are consistent with a previous study which compared the trans-activation and trans-repressive strengths of modern human and mouse PR-A and PR-B and found that the human PR-A was a weaker trans-repressor of PR-B than the mouse PR-A (Wagner, Tong, Emera, and Romero 2012a).

### It is tempting to speculate

Identifying the cause(s) of the relaxed selection intensity acting on PGR in Catarrhine primates the and human stem-lineage is challenging because progesterone signaling has many functions and many traits are phylogenetically associated with Catarrhine primates, any one or none of which may underlie relaxed selection. Given the essential role progesterone plays in female reproduction (Lydon et al. 1995; Conneely et al. 2001; Kubota et al. 2016), however, it is reasonable to conclude that relaxed selection intensity might be related to some aspect of female reproductive biology. Thus, it is interesting to note that Catarrhines have evolved a divergent (and unknown) parturition signal. Unlike nearly every other Eutherian mammal, parturition in Catarrhine primates is not associated with a rapid decline in systemic progesterone concentrations, instead progesterone concentration remains relatively stable throughout pregnancy in Cercopithecidae (‘Old World monkeys’) or continues to increase until parturition in apes (Ratajczak et al. 2010). These data suggest that systematically high progesterone levels throughout pregnancy may have reduced the strength of purifying selection acting on *PGR* in Catarrhines, perhaps because these continually high concentrations compensated for the fixation of deleterious amino acid substitutions.

A more interesting question is whether the enhanced trans-activation ability of PR-B and the weakened trans-repressive strength of PR-A in the human lineage has functional consequences? In the absence of systemic progesterone withdrawal as a mechanism to induce parturition, it has been proposed that progesterone is functionally withdrawn in Catarrhine primates (Csapo 1956; Csapo 1965). Many mechanisms have been proposed to underlie functional progesterone withdrawal, including local breakdown of progesterone in the endometrium and myometrium (Romero et al. 1988; Mazor et al. 1994), sequestration of progesterone in the plasma by carrier proteins (Burton and Westphal 1972; Popp et al. 1978; Heap et al. 1981; Perrot-Applanat and David-Ferreira 1982; Benassayag et al. 2001), a decrease in the levels of PR coactivators at term (Condon et al. 2003), functional estrogen activation (Haluska et al. 1990), and inflammation resulting in NF-κB-mediated PR repression (Allport et al.). A particularly popular hypothesis is that shifts in the abundance of PR-A and PR-B induce functional progesterone withdrawal – when PR-B is the dominant isoform progesterone signaling is active, whereas when PR-A is the dominant isoform progesterone signaling through PR-B is inhibited (Haluska et al., 2002; Merlino et al., 2007; Nadeem et al., 2016; Zakar and Hertelendy, 2007). Consistent with the isoform switching mechanism, the PR-A/PR-B ratio in the myometrium transitions towards the end of pregnancy with PR-A increasing in abundance until parturition in humans (Merlino et al. 2007), macaques (Haluska et al. 2002), and mouse (Zeng et al. 2008). For example, PR-A/PR-B protein ratio in the human myometrium has been reported to be 0.5 (a PR-B dominant state) at 30-week gestation, 1.0 at term prior to the onset of labor, and 3.0 at term during labor (a PR-A dominant state) (Merlino et al. 2007).

While isoform switching is ancestral in primates and rodents and thus did not evolve to initiate parturition, there is evidence supporting its role in the initiation of human labor (Pieber et al. 2001; Mesiano et al. 2002; Mesiano 2004; Merlino et al. 2007; Mittal et al. 2010; Brubaker et al. 2016; Nadeem et al. 2016; Peters et al. 2017; Patel et al. 2018). Furthermore, the role of progesterone in the maintenance of pregnancy up to term is supported by studies showing that disruption of progesterone signaling by the PR-A/PR-B antagonist RU486 at any stage of pregnancy results in myometrial contractions and the initiation of labor in rodents (Dudley et al. 1996; Yang et al. 2000; Shynlova et al. 2013) and primates (Haluska et al., 1990), including humans (Maria et al. 1988). Our observation that the human PR-A is a weaker trans-repressor of PR-B suggests that if isoform switching plays a role in the initiation or progression of parturition, this function may have changed in the human lineage. For example, if repression of PR-B mediated progesterone signaling by PR-A is important for initiating parturition at term, as proposed by the isoform switching hypothesis (Nadeem et al. 2016), then human PR-A may be less effective at inhibiting progesterone signaling and initiating parturition than PR-A of other species (**Figure 4**).

**Figure 4.**
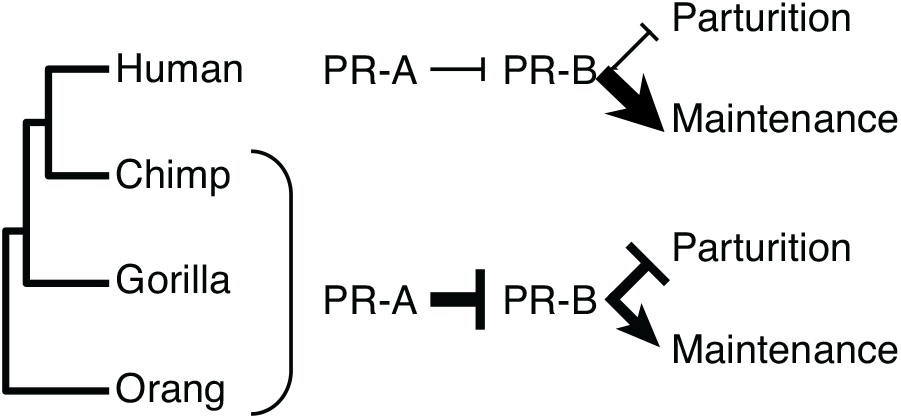
Model of progesterone receptor action under the isoform switching hypothesis. Progesterone signaling through PR-B promotes the maintenance of pregnancy and inhibits parturition, while PR-A induces the cessation of pregnancy and parturition by repressing the actions of PR-B. PR-B evolved a stronger trans-activation ability in the human lineage, while the ability of PR-A to trans-repress PR-B is weaker in the human lineage, suggesting the pregnancy maintenance functions of PR-B are stronger and the parturition inhibiting functions of PR-A are weaker in the human lineage than other species.

### Conclusions, caveats, and limitations

We have shown that *PGR* evolved rapidly in the human stem-lineage, likely reflecting a continuing trend of relaxed purifying selection in Catarrhine primates, rather than positive selection. Regardless of the inference of positive or relaxed selection, we found that the AncHuman PR-A and PR-B diverged in their ability to trans-activate/trans-repress luciferase expression from the dPRL-332 promoter in mouse embryonic fibroblasts (MEFs). Potential limitations of our study include the use of the dPRL-332 promoter and MEFs instead of progesterone-responsive cell types. However, previous studies have shown that the functions of PR-A and PR-B are neither species, cell type, nor promoter-specific and are similar in transiently transfected cells, as well as in cells with a stably integrated reporter gene (Vegeto et al. 1993; Giangrande et al. 2000; Wagner, Tong, Emera, and Romero 2012b) (Wen et al. 1994). Therefore, it is likely that our results will be consistent in other cell types and promoter contexts. Finally, our observations suggest that the regulation of progesterone signaling by PR-A and PR-B might have diverged in the human lineage, meaning that animal models of progesterone signaling may not adequately recapitulate human biology. Thus, the development of organoid models of the human maternal-fetal interface (Rinehart et al. 1988; Boretto et al. 2017; Turco et al. 2017; Turco et al. 2018; Marinić and Lynch 2019) will be important to understand human-specific aspects of progesterone signaling.

## Methods

### Identification of *PGRs* and evolutionary analyses

We identified *PGRs* from BLAST and BLAT searches of assembled and unassembled genomes. Short reads identified from unassembled genomes were assembled with the Geneious ‘map to reference’ option. *PGRs* were aligned with MAFFT (v7) using the auto (FFT-NS-1, FFT-NS-2, FFT-NS-i or L-INS-i) alignment option, 200PAM/k=2 nucleotide scoring matrix, and 1.53 gap opening penalty. Alignment uncertainty was assessed by using GUIDANCE2, which constructed 400 alternative tree topologies by bootstrapping the multiple sequence alignment (MSA) generated by MAFFT (with the best fitting alignment algorithm, FFT-NS-i). Each bootstrap tree is used as a guide tree to re-align the original sequences, and alignment uncertainty inferred from differences in alignment among the bootstrap replicates. The overall GUIDANCE2 alignment score (0.988) was high, indicating MSA uncertainty was low. To ensure that poorly aligned regions of the MSA did not adversely affect downstream analyses, residues with an alignment quality less than 0.67 were masked to N.

We first inferred the best fitting model of nucleotide substitution (012310) using the Datamonkey webserver (Delport et al. 2010). To test if the *d*_*N*_/*d*_*S*_ ratio in the human stem-lineage was significantly different than in other lineages, we compared a model in which the *d*_*N*_/*d*_*S*_ ratio in the human stem-lineage was estimated separately from all other lineages (two rate model) to model with a single *d*_*N*_/*d*_*S*_ ratio for all lineages. This one rate and two rate comparison is implemented in the ‘TestBranchDNDS.bf’ module of HyPhy (2.22) (Pond et al. 2005). Based on exploratory analyses, we allowed for variable *d*_*N*_ and *d*_*S*_ across sites with 3 rate classes. aBSREL (Smith et al. 2015), BUSTED (Murrell et al. 2015), FEL (Pond and Frost 2005), SLAC (Pond and Frost 2005), FUBAR (Murrell et al. 2013), MEME (Murrell et al. 2012), and RELAX (Wertheim et al. 2015) were run using the Datamonkey webserver (Delport et al. 2010), whereas RELAX-scan was run locally using the developer version of HyPhy. We used the Datamonkey webserver and the best fitting model of nucleotide substitution (012310) to infer ancestral sequences. Ancestral human (AncHuman) and ancestral human-chimpanzee (AncHominini) genes were inferred with 1.0 Bayesian Posterior Probabilities. Phylogenetic tests of adaptive evolution (particularly branch-site methods) assume that nucleotide substitutions occur singly and independently. However, multinucleotide mutations (simultaneous mutations at two or three codon positions) can be common and cause false inferences of lineage-specific positive selection (Venkat et al. 2018). We note that human-specific amino acid substitutions result from single nucleotide mutations, G159R (GGG->CGG), A172V (GCT->GTT), S192A (TCC->GCC), G201E (GGG->GAG), P216A (CCC->GCC), S347C (TCT->TGT), P430T (CCC->ACC), and I662V (ATT->GTT), thus these results cannot be explained by the multinucleotide mutations bias.

### Cell culture and luciferase assay

We used mouse embryonic fibroblasts (Wt MEFs, ATCC CRL-2291) to assay PR trans-activation ability, to ensure that difference in trans-activation between ancestral PR proteins was not because of trans-acting factors in human cells interacting more efficiently with AncHuman PR than AncHominini PR or the reverse. MEFs also do not express endogenous *Pgr*, ensuring that the luciferase expression from the dPRL-332 promoter was not affected by endogenous PR levels. MEFs were grown in maintenance medium containing Phenol Red-free DMEM [Gibco] with 10% charcoal-stripped FBS [Gibco], 1x insulin-transferrin-selenium [ITS; Gibco], 1% L-glutamine [Gibco] and 1% sodium pyruvate [Gibco]. 8000 cells were plated per well of a 96-well plate and 18 hours later cells in 80μl of Opti-MEM [Gibco] were transfected with 100ng of one or each of the PR plasmids, 100ng of dPRL-332 vector and 10ng of pRL-null vector with 0.1μl of PLUS Reagent and 0.25μl of Lipofectamine LTX [Invitrogen] in 20μl Opti-MEM. As a negative control, 100ng of dPRL-332 vector, 100ng of pcDNA3.1/V5-HisA vector and 10ng pRL-null vector were used. Cells were incubated with the transfection mixture for 6 hours. Then, cells were washed with DPBS [Gibco] and incubated in the maintenance medium overnight. The next day, decidualization medium (Phenol Red DMEM+GlutaMAX [Gibco] with 2% FBS [Gibco], 1% sodium pyruvate, 0.5mM 8-Br-cAMP [Sigma] and 1μM medroxyprogesterone acetate [MPA; Sigma]) was added and cells were incubated for 48 hours. Upon completion of the treatment, cells were washed with DPBS and incubated for 15 minutes in 1x Passive Lysis Buffer from Dual Luciferase Reporter Assay kit [Promega] with shaking. Luciferase and Renilla activities were measured on Glomax multi+ detection system [Promega] using Luciferin Reagent and Stop&Glow as described in the manufacturer’s protocol. Experiment was repeated 4 times, each condition and control having 12 replicates. We standardized Luciferase activity values to Renilla activity values, and treatment to negative control. Effect size estimates (expressed as log mean difference between groups) and *P*-values were calculated using the DABEST package (Data Analysis with Bootstrap ESTimation) in R (Ho et al. 2019). 5000 bootstrap samples were taken and the confidence interval is bias-corrected and accelerated. The *P*-value(s) reported are the likelihood(s) of observing the effect size(s), if the null hypothesis of zero difference is true.

## References

Csapo AI. 1956. Progesterone block. Am. J. Anat. 98:273–291.

Csapo AI. 1965. The effect of progesterone on the human uterus. Proc. Natl. Acad. Sci. U.S.A. 54:1069–1076.

Abdel-Hafiz H, Takimoto GS, Tung L, Horwitz KB. 2002. The inhibitory function in human progesterone receptor N termini binds SUMO-1 protein to regulate autoinhibition and transrepression. J. Biol. Chem. 277:33950–33956.

Allport VC, Pieber D, human DSM, 2001. Human labour is associated with nuclear factor-κB activity which mediates cyclo-oxygenase-2 expression and is involved with the “functional progesterone withdrawal.” Mol Hum Reprod. 2001 Jun;7(6):581–6.

Arbeitman MN, Furlong EEM, Imam F, Johnson E, Null BH, Baker BS, Krasnow MA, Scott MP, Davis RW, White KP. 2002. Gene Expression During the Life Cycle of Drosophila melanogaster. Science 297:2270–2275.

Arbiza L, Dopazo J, Dopazo H. 2006. Positive selection, relaxation, and acceleration in the evolution of the human and chimp genome. PLoS Comput. Biol. 2:e38.

Bakewell MA, Shi P, Zhang J. 2007. More genes underwent positive selection in chimpanzee evolution than in human evolution. Proc. Natl. Acad. Sci. U.S.A. 104:7489–7494.

Behringer RR, Eakin GS, Renfree MB. 2006. Mammalian diversity: gametes, embryos and reproduction. Reprod. Fertil. Dev. 18:99–107.

Benassayag C, Souski I, Mignot TM, Robert B, Hassid J, Duc-Goiran P, Mondon F, Rebourcet R, Dehennin L, Nunez EA, et al. 2001. Corticosteroid-binding globulin status at the fetomaternal interface during human term pregnancy. Biol. Reprod. 64:812–821.

Benshoof L, Thornhill R. 1979. The evolution of monogamy and concealed ovulation in humans. Journal of Social and Biological Structures 2:95–106.

Boretto M, Cox B, Noben M, Hendriks N, Fassbender A, Roose H, Amant F, Timmerman D, Tomassetti C, Vanhie A, et al. 2017. Development of organoids from mouse and human endometrium showing endometrial epithelium physiology and long-term expandability. Development 144:1775–1786.

Bradshaw FJ, Bradshaw D. 2011. Progesterone and reproduction in marsupials: a review. General and Comparative Endocrinology 170:18–40.

Burley N. 1979. The Evolution of Concealed Ovulation. The American Naturalist 114:835–858.

Burton RM, Westphal U. 1972. Steroid hormone-binding proteins in blood plasma. Metabolism 21:253–276.

Chen C, Opazo JC, Erez O, Uddin M, Santolaya-Forgas J, Goodman M, Grossman LI, Romero R, Wildman DE. 2008. The human progesterone receptor shows evidence of adaptive evolution associated with its ability to act as a transcription factor. Mol. Phylogenet. Evol. 47:637–649.

Christian M, Pohnke Y, Kempf R, Gellersen B, Brosens JJ. 2002. Functional Association of PR and CCAAT/Enhancer-Binding Protein {beta} Isoforms: Promoter-Dependent Cooperation between PR-B and Liver-Enriched Inhibitory Protein, or Liver-Enriched Activatory Protein and PR-A in Human Endometrial Stromal Cells. Mol. Endocrinol. 16:141–154.

Christian M, Zhang X, Schneider-Merck T, Unterman TG, Gellersen B, White JO, Brosens JJ. 2002. Cyclic AMP-induced Forkhead Transcription Factor, FKHR, Cooperates with CCAAT/Enhancer-binding Protein Œ≤ in Differentiating Human Endometrial Stromal Cells. J. Biol. Chem. 277:20825–20832.

Clark AG, Glanowski S, Nielsen R, Thomas PD, Kejariwal A, Todd MA, Tanenbaum DM, Civello D, Lu F, Murphy B, et al. 2003. Inferring nonneutral evolution from human-chimp-mouse orthologous gene trios. Science 302:1960–1963.

Condon JC, Jeyasuria P, Faust JM, Wilson JW, Mendelson CR. 2003. A decline in the levels of progesterone receptor coactivators in the pregnant uterus at term may antagonize progesterone receptor function and contribute to the initiation of parturition. Proc. Natl. Acad. Sci. U.S.A. 100:9518–9523.

Conneely OM, Mulac-Jericevic B, Lydon JP, De Mayo FJ. 2001. Reproductive functions of the progesterone receptor isoforms: lessons from knock-out mice. Mol Cell Endocrinol 179:97–103.

Delport W, Poon AFY, Frost SDW, Kosakovsky Pond SL. 2010. Datamonkey 2010: a suite of phylogenetic analysis tools for evolutionary biology. Bioinformatics 26:2455–2457.

Dudley DJ, Branch DW, Edwin SS, Mitchell MD. 1996. Induction of preterm birth in mice by RU486. Biol. Reprod. 55:992–995.

Emera D, Romero R, Wagner G. 2012. The evolution of menstruation: A new model for genetic assimilation. Bioessays 34:26–35.

George RD, McVicker G, Diederich R, Ng SB, MacKenzie AP, Swanson WJ, Shendure J, Thomas JH. 2011. Trans genomic capture and sequencing of primate exomes reveals new targets of positive selection. Genome Res. 21:1686–1694.

Giangrande PH, Kimbrel EA, Edwards DP, McDonnell DP. 2000. The opposing transcriptional activities of the two isoforms of the human progesterone receptor are due to differential cofactor binding. Mol. Cell. Biol. 20:3102–3115.

Graham CE. 1981. Reproductive Biology of the Great Apes: Comparative and Biomedical Perspectives. Academic Press

Haluska GJ, Wells TR, Hirst JJ, Brenner RM, Sadowsky DW, Novy MJ. 2002. Progesterone receptor localization and isoforms in myometrium, decidua, and fetal membranes from rhesus macaques: evidence for functional progesterone withdrawal at parturition. J. Soc. Gynecol. Investig. 9:125–136.

Haluska GJ, West NB, Novy MJ, Brenner RM. 1990. Uterine estrogen receptors are increased by RU486 in late pregnant rhesus macaques but not after spontaneous labor. J. Clin. Endocrinol. Metab. 70:181–186.

Hamilton ME. 1984. Revising Evolutionary Narratives: A Consideration of Alternative Assumptions about Sexual Selection and Competition for Mates. American Anthropologist 86:651–662.

Heap RB, Ackland N, Weir BJ. 1981. Progesterone-binding proteins in plasma of guinea-pigs and other hystricomorph rodents. J. Reprod. Fertil. 63:477–489.

Ho J, Tumkaya T, Aryal S, Choi H, Claridge-Chang A. 2019. Moving beyond P values: Everyday data analysis with estimation plots. bioRxiv 2:377978.

Huse B, Verca SB, Matthey P, Rusconi S. 1998. Definition of a negative modulation domain in the human progesterone receptor. Mol. Endocrinol. 12:1334–1342.

Jiang Y, Hu Y, Zhao J, Zhen X, Yan G, Sun H. 2011. The orphan nuclear receptor Nur77 regulates decidual prolactin expression in human endometrial stromal cells. Biochem Biophys Res Commun 404:628–633.

Kastner P, Krust A, Turcotte B, Stropp U, Tora L, Gronemeyer H, Chambon P. 1990. Two distinct estrogen-regulated promoters generate transcripts encoding the two functionally different human progesterone receptor forms A and B. EMBO J. 9:1603–1614.

Kaya HS, Hantak AM, Stubbs LJ, Taylor RN, Bagchi IC, Bagchi MK. 2015. Roles of progesterone receptor A and B isoforms during human endometrial decidualization. Mol. Endocrinol. 29:882–895.

Kin K, Maziarz J, Wagner GP. 2014. Immunohistological Study of the Endometrial Stromal Fibroblasts in the Opossum, Monodelphis domestica: Evidence for Homology with Eutherian Stromal Fibroblasts. Biol Reprod. 2014 May;90(5):111. doi: 10.1095/biolreprod.113.115139.

Kircher M, Witten DM, Jain P, O’Roak BJ, Cooper GM, Shendure J. 2014. A general framework for estimating the relative pathogenicity of human genetic variants. Nat. Genet. 46:310–315.

Kosakovsky Pond SL, Murrell B, Fourment M, Frost SDW, Delport W, Scheffler K. 2011. A random effects branch-site model for detecting episodic diversifying selection. Mol Biol Evol. 2011 Nov;28(11):3033–43. doi: 10.1093/molbev/msr125

Kosiol C, Vinar T, da Fonseca RR, Hubisz MJ, Bustamante CD, Nielsen R, Siepel A. 2008. Patterns of Positive Selection in Six Mammalian Genomes. Schierup MH, editor. PLoS Genet. 4:e1000144.

Kubota K, Cui W, Dhakal P, Wolfe MW, Rumi MAK, Vivian JL, Roby KF, Soares MJ. 2016. Rethinking progesterone regulation of female reproductive cyclicity. Proc. Natl. Acad. Sci. U.S.A. 113:4212–4217.

Lombardi J. 1998. Comparative Verbebrate Reproduction. 1st ed. Boston: Kluwer Academic Publishers

Lydon JP, DeMayo FJ, Funk CR, Mani SK, Hughes AR, Montgomery CA, Shyamala G, Conneely OM, O’Malley BW. 1995. Mice lacking progesterone receptor exhibit pleiotropic reproductive abnormalities. Genes Dev. 9:2266–2278.

Lynch VJ, Brayer K, Gellersen B, Wagner GP. 2009. HoxA-11 and FOXO1A cooperate to regulate decidual prolactin expression: towards inferring the core transcriptional regulators of decidual genes. PLoS ONE 4:e6845.

Maria B, Stampf F, Goepp A, Ulmann A. 1988. Termination of early pregnancy by a single dose of mifepristone (RU 486), a progesterone antagonist. Eur. J. Obstet. Gynecol. Reprod. Biol. 28:249–255.

Marinić M, Lynch VJ. 2019. Derivation of endometrial gland organoids from term post-partum placenta. bioRxiv:753780.

Mascia-Lees FE, Relethford JH, Sorger T. 1986. Evolutionary Perspectives on Permanent Breast Enlargement in Human Females. American Anthropologist 88:423–428.

Mazor M, Hershkovitz R, Chaim W, Levy J, Sharony Y, Leiberman JR, Glezerman M. 1994. Human preterm birth is associated with systemic and local changes in progesterone/17 beta-estradiol ratios. Am. J. Obstet. Gynecol. 171:231–236.

Meiklejohn CD, Parsch J, Ranz JM, Hartl DL. 2003. Rapid evolution of male-biased gene expression in Drosophila. Proc. Natl. Acad. Sci. U.S.A. 100:9894–9899.

Merlino AA, Welsh TN, Tan H, Yi LJ, Cannon V, Mercer BM, Mesiano S. 2007. Nuclear progesterone receptors in the human pregnancy myometrium: evidence that parturition involves functional progesterone withdrawal mediated by increased expression of progesterone receptor-A. J. Clin. Endocrinol. Metab. 92:1927–1933.

Mess A, Carter AM. 2006. Evolutionary transformations of fetal membrane characters in Eutheria with special reference to Afrotheria. J. Exp. Zool. B Mol. Dev. Evol. 306B:140–163.

Murrell B, Moola S, Mabona A, Weighill T, Sheward D, Kosakovsky Pond SL, Scheffler K. 2013. FUBAR: a fast, unconstrained bayesian approximation for inferring selection. Mol. Biol. Evol. 30:1196–1205.

Murrell B, Weaver S, Smith MD, Wertheim JO, Murrell S, Aylward A, Eren K, Pollner T, Martin DP, Smith DM, et al. 2015. Gene-wide identification of episodic selection. Mol. Biol. Evol. 32:1365–1371.

Murrell B, Wertheim JO, Moola S, Weighill T, Scheffler K, Kosakovsky Pond SL. 2012. Detecting individual sites subject to episodic diversifying selection. PLoS Genet. 8:e1002764.

Nadeem L, Shynlova O, Matysiak-Zablocki E, Mesiano S, Dong X, Lye S. 2016. Molecular evidence of functional progesterone withdrawal in human myometrium. Nat Commun 7:11565–11569.

Nielsen R, Bustamante C, Clark AG, Glanowski S, Sackton TB, Hubisz MJ, Fledel-Alon A, Tanenbaum DM, Civello D, White TJ, et al. 2005. A scan for positively selected genes in the genomes of humans and chimpanzees. PLoS Biol. 3:e170.

Parisi M, Nuttall R, Naiman D, Bouffard G, Malley J, Andrews J, Eastman S, Oliver B. 2003. Paucity of Genes on the Drosophila X Chromosome Showing Male-Biased Expression. Science 299:697–700.

Perrot-Applanat M, David-Ferreira JF. 1982. Immunocytochemical localization of progesterone-binding protein (PBP) in guinea-pig placental tissue. Cell Tissue Res. 223:627–639.

Phillips JB, Abbot P, Rokas A. 2015. Is preterm birth a human-specific syndrome? Evol Med Public Health 2015:136–148.

Pohnke Y, Kempf R, Gellersen B. 1999. CCAAT/Enhancer-binding Proteins Are Mediators in the Protein Kinase A-dependent Activation of the Decidual Prolactin Promoter. J. Biol. Chem. 274:24808–24818.

Kosakovsky Pond SL, Frost S. 2005. Not so different after all: a comparison of methods for detecting amino acid sites under selection. Mol Biol Evol. 2005 May;22(5):1208–22.

Kosakovsky Pond SL, Frost SDW, Muse SV. 2005. HyPhy: hypothesis testing using phylogenies. Bioinformatics 21:676–679.

Popp RA, Foresman KR, Wise LD, Daniel JC. 1978. Amino acid sequence of a progesterone-binding protein. Proc. Natl. Acad. Sci. U.S.A. 75:5516–5519.

Ratajczak CK, Fay JC, Muglia LJ. 2010. Preventing preterm birth: the past limitations and new potential of animal models. Dis Model Mech 3:407–414.

Renfree M, Shaw G. 2001. Reproduction in Monotremes and Marsupials. In: eLS. John Wiley & Sons, Ltd.

Rentzsch P, Witten D, Cooper GM, Shendure J, Kircher M. 2019. CADD: predicting the deleteriousness of variants throughout the human genome. Nucleic Acids Res. 47:D886–D894.

Rinehart CA, Lyn-Cook BD, Kaufman DG. 1988. Gland formation from human endometrial epithelial cells in vitro. In Vitro Cell. Dev. Biol. 24:1037–1041.

Romero R, Scoccia B, Mazor M, Wu YK, Benveniste R. 1988. Evidence for a local change in the progesterone/estrogen ratio in human parturition at term. Am. J. Obstet. Gynecol. 159:657–660.

Shynlova O, Nedd-Roderique T, Li Y, Dorogin A, Lye SJ. 2013. Myometrial immune cells contribute to term parturition, preterm labour and post-partum involution in mice. J. Cell. Mol. Med. 17:90–102.

Smith MD, Wertheim JO, Weaver S, Murrell B, Scheffler K, Kosakovsky Pond SL. 2015. Less is more: an adaptive branch-site random effects model for efficient detection of episodic diversifying selection. Mol. Biol. Evol. 32:1342–1353.

Strassmann BI. 2015. The Evolution of Endometrial Cycles and Menstruation. The Quarterly Review of Biology 71:181–220.

Turco MY, Gardner L, Hughes J, Cindrova-Davies T, Gomez MJ, Farrell L, Hollinshead M, Marsh SGE, Brosens JJ, Critchley HO, et al. 2017. Long-term, hormone-responsive organoid cultures of human endometrium in a chemically defined medium. Nat. Cell Biol. 19:568–577.

Turco MY, Gardner L, Kay RG, Hamilton RS, Prater M, Hollinshead MS, McWhinnie A, Esposito L, Fernando R, Skelton H, et al. 2018. Trophoblast organoids as a model for maternal–fetal interactions during human placentation. Nature 564:263–267.

van der Lee R, Wiel L, van Dam TJP, Huynen MA. 2017. Genome-scale detection of positive selection in nine primates predicts human-virus evolutionary conflicts. Nucleic Acids Res. 45:10634–10648.

Vegeto E, Shahbaz MM, Wen DX, Goldman ME, O’Malley BW, McDonnell DP. 1993. Human progesterone receptor A form is a cell- and promoter-specific repressor of human progesterone receptor B function. Mol. Endocrinol. 7:1244–1255.

Venkat A, Hahn MW, Thornton JW. 2018. Multinucleotide mutations cause false inferences of lineage-specific positive selection. Nat Ecol Evol 2:1280–1288.

Wagner GP, Tong Y, Emera D, Romero R. 2012a. An evolutionary test of the isoform switching hypothesis of functional progesterone withdrawal for parturition: humans have a weaker repressive effect of PR-A than mice. J Perinat Med 40:345–351.

Wen DX, Xu YF, Mais DE, Goldman ME, McDonnell DP. 1994. The A and B isoforms of the human progesterone receptor operate through distinct signaling pathways within target cells. Mol. Cell. Biol. 14:8356–8364.

Wertheim JO, Murrell B, Smith MD, Kosakovsky Pond SL, Scheffler K. 2015. RELAX: Detecting Relaxed Selection in a Phylogenetic Framework. Mol. Biol. Evol. 32:820–832.

Wetendorf M, DeMayo FJ. 2012. The progesterone receptor regulates implantation, decidualization, and glandular development via a complex paracrine signaling network. Mol Cell Endocrinol 357:108–118.

Yang B, Zhou HJ, He QJ, Fang RY. 2000. Termination of early pregnancy in the mouse, rat and hamster with DL111-IT and RU486. Contraception 62:211–216.

Zeng Z, Velarde MC, Simmen FA, Simmen RCM. 2008. Delayed parturition and altered myometrial progesterone receptor isoform A expression in mice null for Krüppel-like factor 9. Biol. Reprod. 78:1029–1037.

